# Sauna-like conditions or menthol treatment reduce tau phosphorylation through mild hyperthermia

**DOI:** 10.1101/2021.01.27.428475

**Authors:** Isabelle Guisle, Séréna Pétry, Françoise Morin, Rémi Kérauden, Robert A. Whittington, Frédéric Calon, Sébastien S. Hébert, Emmanuel Planel

## Abstract

In Alzheimer’s disease (AD), hyper-phosphorylation and aggregation of tau correlates with clinical progression and represents a valid therapeutic target. A recent 20-year prospective study revealed an association between moderate to high frequency of Finnish sauna bathing and a lower incidence of dementia and AD, but the molecular mechanisms underlying these benefits remain uncertain. Here, we tested the hypothesis that sauna-like conditions could lower tau phosphorylation by increasing body temperature. We observed a decrease in tau phosphorylation in wild-type and hTau mice as well as in neuron-like cells when exposed to higher temperatures. These effects were correlated with specific changes in phosphatase and kinase activities, but not with inflammatory or heat-shock responses. We also used a drug strategy to promote thermogenesis. Topical application of menthol led to a sustained increase in body temperature in hTau mice concomitant with a significant decrease in tau phosphorylation. Our results suggest that sauna-like conditions or menthol treatment could lower tau pathology through mild hyperthermia, and may provide promising therapeutic strategies for AD and other tauopathies.

## Introduction

Tau protein is a microtubule-associated protein that binds to and stabilizes microtubules (Weingarten et al. 1975). Under physiological conditions, phosphorylation regulates tau binding to microtubules and many of its other funtions (Guo et al. 2017). However, abnormal hyperphosphorylation of tau has been shown to lead to agregation and neurofibrillary tangle formation (Alonso et al. 1996), a key feature of several diseases known as tauopathies, the most common being Alzheimer’s disease (AD) (Avila et al. 2004). The neurodegeneration and clinical progression of AD are correlated with tau pathology (Braak and Del Tredici 2016), making tau phosphorylation a valid target for the treatment of AD and other tauopathies. However, and despite many efforts, no treatment currently exists for tauopathies (Rosler et al. 2020).

Recently, a 20-year longitudinal study of more than 2000 Finnish revealed that sauna bathing between 4 to 7 times a week lowered the risk of developing AD when compared to once a week (Laukkanen T.S.K 2016). Sauna bathing is associated with well-being and relaxation as well as reduction in systemic blood pressure, improvement of endothelial function, and reductions in oxidative stress and inflammation (Laukkanen et al. 2018). Therefore, it was proposed that such responses could underlie the beneficial effects of sauna bathing on the risk of AD. However, the effects of sauna on AD biomarkers were not investigated (Laukkanen T.S.K 2016). Interestingly, another consequence of sauna is to induce mild hyperthermia (Sohar et al. 1976). The temperature of a Finnish sauna is usually between 80 to 100°C, with humidity varying between 10 to 40%. Under these conditions, 20 min is sufficient to raise body temperature to approximately 38-39°C (Sohar, Shoenfeld et al. 1976).

In animal models, hypothermia consistently induces tau hyper-phosphorylation (Planel et al. 2004, Whittington et al. 2010, Bretteville et al. 2012, Vandal et al. 2016, Tournissac et al. 2017). Therefore, we hypothesized that, in contrast to hypothermia, mild hyperthermia could lower tau phosphorylation. Indeed, we observed that mild hyperthermia induced tau de-phosphorylation in both wild-type and hTau mice, a model of AD-like tauopathy. Similar results were observed in neuron like cells expressing human tau.

As sauna bathing is not openly accessible to most, we examined a therapeutic way to increase body temperature by using menthol. Menthol is a known activator of TRPM8 (Transient Receptor Potential Melastatin 8) or CMR1 (Cold and Menthol Receptor 1), a calcium-gated channel triggered by heat/cool exposure (McKemy et al. 2002, Peier et al. 2002). TRPM8 is expressed in somatosensory neurons that innervate peripheral tissues such as the skin and oral cavity, and is the primary molecular transducer of cold somatosensation (hence the “cooling” effect of menthol) (McCoy et al. 2011). Activation of TRPM8 by topical menthol administration promotes thermogenesis and an increase in temperature in mice and humans (Tajino et al. 2007, Gillis et al. 2015). Thus, we hypothesized that heat production induced by menthol treatment could lead to dephosphorylation of tau. Topical application of menthol resulted in transient increase in body temperature and tau dephosphorylation in hTau mice, supporting our hypothesis.

Altogether, our results suggest that sauna-like conditions and menthol treatment can lower tau pathology through mild hyperthermia and may partially explain the beneficial effects of sauna bathing observed by *Laukkanen et al*. These findings open new potential therapeutic treatments that are discussed here.

## Material and methods

### Animals

8 week-old C57BL6 and 12 month-old hTau female mice (Andorfer et al. 2003) were used for heat exposure (n=11-12). The hTau mice (Andorfer, Kress et al. 2003) were obtained by crossing mice that express the six isoforms of human non mutated tau: 8c mice (Duff et al. 2000)) with murine tau knock out mice (Tucker et al. 2001). The founders of our hTau and tau knock out colonies were derived on a C57BL6/J background (B6.Cg-Mapttm1(EGFP)Klt-Tg(MAPT)8cPdav/J, Jackson Laboratories, Bar Harbor, ME, USA). hTau mice accumulate hyperphosphorylated tau and aggregated tau at 6 and 9 month of age, respectively (Andorfer, Kress et al. 2003). The animals were handled according to procedures approved by the “Comité de Protection des Animaux du CHU” under the guidelines of the Canadian Council on Animal Care. Mice were killed by decapitation without anesthesia, as anesthesia can lead to hypothermia-induced tau phosphorylation (Planel et al. 2007).

### Heat treatment

To induce mild hyperthermia (39-40°C rectal temperature), the mice were kept in a ventilated incubator at 42°C for 45 min. Control mice where kept at ambient temperature (23°C). All mice had access to water and food *ad libitum*. Core body temperature was assessed at the end of the incubation, just prior to sacrifice, with a rectal probe (RET-3, Brain Tree Scientific Inc, Braintree, MA, USA).

### Menthol treatment

Vehicle and menthol treated mice were housed in a ventilated incubator at 28°C for three day before the experiment, as previously described (Tajino, Matsumura et al. 2007). Topical application of 10% menthol (weight: volume, in ethanol 100%) or vehicle (ethanol 100%) (W266523, Sigma Aldrich, St. Louis, MO, USA) was done as previously described (Tajino, Matsumura et al. 2007). Briefly, animals were rapidly anesthetized with 3% isoflurane (no more than 2min for each application). A 5cm x 6cm tissue paper (Kimwipe, S-8115, Kimtech, Milton, ON, Canada) was soaked with 600µL of 10% menthol or 100% ethanol and applied to the body of the animals (except the head, limb and tail). The fur of the animals was not shaved. Application of menthol 10% or ethanol was done once in the morning between 10h30 and 11h45. The rectal temperature of the animals was taken 2 hours after exposure to menthol or ethanol, the mice were immediately sacrificed by decapitation and the brains were rapidly removed and dissected (see section tissue extraction). Glycemia was assessed with blood obtained at decapitation using a glucometer with reagent strips (ACCU-CHEK ^®^ Aviva Nano; Roche Diagnostics GmbH, Mannheim, Germany).

### Assessment of body temperature

To evaluate the effect of heat treatment or 10% menthol treatment on body temperature, telemetric probes (Anipill precision 02, BodyCap, Caen, France) were surgically implanted in the abdomen of 21 month-old C57Bl6 male mice under isoflurane anesthesia and recorded at 23 month-old. Temperature was recorded every 5 minutes and all data points were stored for analyses. Heat exposed mice, 10% menthol exposed mice and their respective controls were treated exactly the same manner as described above.

### Cell culture

Human neuroblastoma cells (SH-SY5Y cells) stably expressing human tau 3 repeat isoform 2+3-10- (generous gift from Luc Buée) were used for this experiment. SH-SY5Y 3R-tau cells were cultured as previously described (Delobel et al. 2003) in a 5% CO2 incubator for 24h either at 34°C, at 37°C or at 40°C. At the end of the incubation, cells were rapidly put on ice, the culture medium was removed, and the cells were washed twice with phospho-buffered saline 1x, collected in 100µL of Radioimmunoprecipitation Assay and processed as previously described (Whittington et al. 2011). 10 μg of protein were analyzed as described previously (Petry et al. 2014, Petry et al. 2017).

### Tissue extraction

Brains were immediately removed and tissues were dissected on ice. Tissues were frozen on dry ice and kept at − 80 °C until they were homogenized. The cortices were homogenized without thawing in 5 times volume-weight of RIPA with a mechanical homogenizer (14-261-01, Fisher Scientific, Waltham, MA, USA). The hippocampi were homogenized without thawing by sonication in 150*µ*L of RIPA. Samples were then centrifuged for 20 min at 20,000*g* at 4 °C. The supernatant was recovered, diluted in sample buffer (NuPAGE LDS; Invitrogen, Carlsbad, CA) containing 5% of 2-β-mercapto-ethanol, 1 mM Na_3_VO_4_, 1 mM NaF, 1 mM PMSF, 10 μl/ml of Proteases Inhibitors Cocktail (P8340; Sigma-Aldrich, St. Louis, MO, USA), and heated for 10 min at 95 °C. 10 μg-20µg of protein were analyzed as described previously (Petry, Pelletier et al. 2014, Petry, Nicholls et al. 2017).

### Protein analysis

SDS-PAGE and Western blot analysis were done as previously described (Planel et al. 2001, Gratuze et al. 2017). Briefly, 10µg of brain homogenates were separated on a SDS-10% polyacrylamide gel, transferred onto nitrocellulose membranes (Amersham Biosciences, Pittsburgh, PA, USA). Membranes where then saturated, hybridized with antibodies and revealed as described in (Gratuze, El Khoury et al. 2017). The analysis of tau aggregates was done by sarkosyl extraction, according to a protocol derived from Greenberg and Davis (Greenberg and Davies 1990, Julien et al. 2012). All antibodies used in this study, in addition to their dilution, are listed in **Table 1**. For tau protein immunoblots, the signal was normalized to total tau for all epitopes. For kinases immunoblots, the signal was normalized to the total corresponding protein. Two lanes from representative immunoblots are displayed for each condition. Dividing lines represent areas where lanes from the same blot were removed and the remaining lanes were spliced together. Brightness levels were adjusted as needed.

**Table 1:** Antobodies used in this study (in supplemental data)

| Name | Abbreviation | Epitope/Immunogen | type | host | Provider | Reference | WB dilution |
| --- | --- | --- | --- | --- | --- | --- | --- |
| **tau antibodies** |  |  |  |  |  |  |  |
| anti tau C | TauC (C terminal) | C-terminal part of tau (amino acids 243-441) | polyclonal | Rabbit | Dako Cytomation | A0024 | 1/10000 |
| anti tau 1, clone PC1C6 | Tau-1 | phospho-serine 195, 198, 199 and 202 (recognizes dephosphorylated tau) | monoclonal | Mouse IgG<sub>2a</sub> | Millipore | MAB3420 | 1/10000 |
| anti phospho tau antibody pS404 | pS404 | phospho-serine 404 | polyclonal | Rabbit | Life Technologies | 44758G | 1/1000 |
| anti phospho Tau antibody: clone AT8 | AT8 | phospho-serine 202, phospho-threonine 205 | monoclonal | Mouse IgG1k | Fisher | MN1020 | 1/1000 |
| anti phospho Tau antibody: clone AT180 | AT180 | phospho-threonine 231 | monoclonal | Mouse IgG1k | Fisher | MN1040 | 1/1000 |
| anti phospho tau antibody CP13 | CP13 | phospho-serine 202 | monoclonal | Mouse | Peter Davies | - | 1/1000 |
| anti phospho tau antibody PHF1 | PHF1 | phospho-serine 396, 404 | monoclonal | Mouse | Peter Davies | - | 1/1000 |
| **Kinase Antibodies** |  |  |  |  |  |  |  |
| anti akt protein (anti protein kinase B) | Akt | carboxy-terminal sequence of mouse Akt1, Akt2 and Akt3 proteins | polyclonal | Rabbit | NEB | 9272 | 1/1000 |
| anti phospho Akt | Phospho-Akt | Akt1-2-3 phosphorylated at Ser473 | polyclonal | Rabbit | NEB | 9271 | 1/1000 |
| anti Adenosine monophosphate protein kinase $\alpha$ subunit (AMPKα) | AMPKα | amino-terminal sequence of human AMPKα | monoclonal | Rabbit | NEB | 2603 | 1/1000 |
| anti phospho AMPKα1-2 subunit (clone 40H9) | p-AMPKα | phospho-threonine 172 | monoclonal | Rabbit | NEB | 2535 | 1/1000 |
| anti calcium calmodulin Kinase II alpha subunit | CaMKII(alpha) | mouse CaMKIIα amino-acids 303-478 | monoclonal | Mouse | Santa Cruz | sc-13141 | 1/1000 |
| anti Phospho-CaMKII(a) | p-CaMKII(a) | phospho-threonine 286 | polyclonal | Rabbit | Santa Cruz | sc-12886 | 1/1000 |
| anti glycogen synthase kinase 3 b | pGSK-3b | phospho-serine 9 | polyclonal | Rabbit | NEB | 9336 | 1/1000 |
| anti GSK-3β | GSK-3β | Rat GSK-3β amino acid 1-160 | monoclonal | Mouse IgG1 | BD Biosciences | 610202 | 1/1000 |
| p44/42 Mitogen Activated Protein Kinase/Extracellular signal Regulated Kinase 1/2 | ERK | total p44/42 MAP kinase (Erk1/Erk2) protein | polyclonal | Rabbit | NEB | 9102 | 1/1000 |
| Phospho-p44/42 MAPK (Erk1/2) Antibody | pERK | phospho Threonine 202/phosphoTyrosine 204) | polyclonal | Rabbit | NEB | 9101 | 1/1000 |
| stress-activated protein kinase/Jun-amino-terminal kinase | SAPK/JNK | total JNK1, JNK2 or JNK3 protein | polyclonal | Rabbit | NEB | 9252 | 1/1000 |
| Phospho-SAPK/JNK Antibody | phospho-SAPK/JNK | phospho-threonine 183/phospho-tyrosine 185 | polyclonal | Rabbit | NEB | 9251 | 1/1000 |
| Protein Kinase A a/b catalytic subunit (clone H56) | PKAα/βcat | c terminal amino acids 231-286 of PKAα catalytic subunit from human origin | polyclonal | Rabbit | Santa Cruz | sc-98951 | 1/1000 |
| phospho PKAa/b | p-PKAα/$\beta$/$\gamma$ cat | phospho Threonine 198 | polyclonal | Rabbit | Santa Cruz | sc-32968 | 1/1000 |
| **Phosphatase Antibodies** |  |  |  |  |  |  |  |
| Anti-protein phosphatase 2A antibody, C subunit, clone 7A6 | PP2A, C subunit | amino acids 302-309 of human PP2A catalytic subunit | monoclonal | Mouse | Millipore | 05-545 | 1/1000 |
| anti demethylated PP2A C subunit | Demethylated-PP2A-C | unmethylated C-terminal region of the PP2A-C subunit | monoclonal | Mouse | Santa Cruz | sc-13601 | 1/1000 |
| **control Antibodies** |  |  |  |  |  |  |  |
| anti beta actin clone AC-74 | Beta-Actin | synthetic $\beta$ cytoplasmic actin N-terminal peptide | monoclonal | Mouse | Sigma | A2228 | 1/10000 |
| anti Glyceraldehyde-3-phosphate dehydrogenase antibody clone 6C5 | GAPDH | GAPDH | monoclonal | Mouse | Millipore | MAB374 | 1/10000 |
| **Secondary antibodies** |  |  |  |  |  |  |  |
| peroxidase conjugated affinipure goat anti mouse IgG | anti mouse | Mouse IgG | - | Goat | Jackson Immunoresearch | - | 1/5000 |
| peroxidase conjugated affinipure goat anti rabbit IgG | anti rabbit | Rabbit IgG | - | Goat | Jackson Immunoresearch | - | 1/5000 |
| Goat anti mouse light chain antibody, horse radish peroxidase conjugate | anti Light Chain | reactive with native primary antibodies light chain | - | Goat | Millipore | - | 1/5000 |
| **Other Antibodies** |  |  |  |  |  |  |  |
| Toll Like Receptor 2 antibody | TLR2 | Extracellular domain of human TLR2. | polyclonal | Rabbit | Millipore | 06-1119 | 1/1000 |
| Glial Fibrillary Acidic Protein | GFAP | GFAP | monoclonal | Mouse | Sigma | G3893 | 1/1000 |
| Heat shock protein 70 | Hsp70 | Human Hsp70 aa. 429-640 | monoclonal | Mouse | BD Biosciences | 610607 | 1/1000 |

### Analysis of aggregated tau

Tau aggregates were extracted according to previously published protocols derived from Briefly, the RIPA supernatant was adjusted to 1% sarkosyl (*N*-lauroylsarcosine), incubated for 60 min at 37°C with constant shaking, and centrifuged at 100,000×*g* for 1 h at 20 °C. The pellet containing sarkosyl-insoluble aggregated tau was resuspended and diluted in sample buffer (NuPAGE LDS) containing 5% of 2-β -mercapto-ethanol, 1 mM Na_3_VO_4_, 1 mM NaF, 1 mM PMSF, 10 µl/ml of proteases inhibitors cocktail (P8340, Sigma-Aldrich), sonicated and boiled for 5 min, and then stored at −20 °C.

### Protein Phosphatase 2 A (PP2A) activity assay

A 2 months old C57BL6 mouse was killed by decapitation without anesthesia. The brain was removed, rapidly dissected on ice and frozen at −80°C until they were used. The frozen cortex was homogenized according to the manufacturer’s of the Duo set IC active PP2A kit (reference DYC3309, R&D systems, Minneapolis, USA) and has been previously described (Gratuze et al. 2017). The lysate was processed at 2 µg/µL in triplicates (n=3) with the same kit at 37°C, 40°C and 42°C.

### Tau kinase activity assay

Brain tau kinase assay was done according to our previous protocol (Planel, Miyasaka et al. 2004) with some modifications. 1µg of full length recombinant tau (reference T1001-2, R Peptide, Watkinsville, GA, USA) was prepared in sample buffer containing 5µg of RIPA cortex lysate. This reaction was incubated for 60 minutes in triplicates for each temperature (37°C, 40°C and 42°C) and terminated by adding one volume of sample buffer (NuPAGE LDS; Invitrogen, Carlsbad, CA) containing 5% of 2-β-mercapto-ethanol, 1 mM Na_3_VO_4_, 1 mM NaF, 1 mM PMSF, 10 μl/ml of Proteases Inhibitors Cocktail (P8340; Sigma-Aldrich, St. Louis, MO, USA), and heated for 10 min at 95 °C. A negative control without ATP was processed in parallel following the same procedure in duplicate for each temperature. Samples were analyzed by SDS-PAGE and western blotting as described above. Immunoblot was analyzed using anti tau PS404 antibody (see section antibodies) since this epitope is phosphorylated by most of the major tau of kinases (Planel 2002). The signal was normalized to total tau. Immunoreactive bands were visualized using ImageQuant LAS 4000 imaging system (Fujifilm USA, Valhalla, NY, USA), and densitometric analyses were performed with Image Gauge analysis software (Fujifilm, Valhalla, NY, USA). Images levels were adjusted.

### Statistical analysis

Non parametric data were analyzed using a Man Whitney test. For normally distributed data, an unpaired t-test with Wech’s correction was applied. Figure 5 was analyzed by Kruskall Wallis test followed by Dunn’s post hoc test for pair-wise sample comparison. All data calculations were performed using GraphPad Prism software 5.0 (GraphPad Software, La Jolla, CA, USA). Significant results were given for p<0.05. For all figures, data are given as mean ± Standard Deviation.

**Figure 1:**
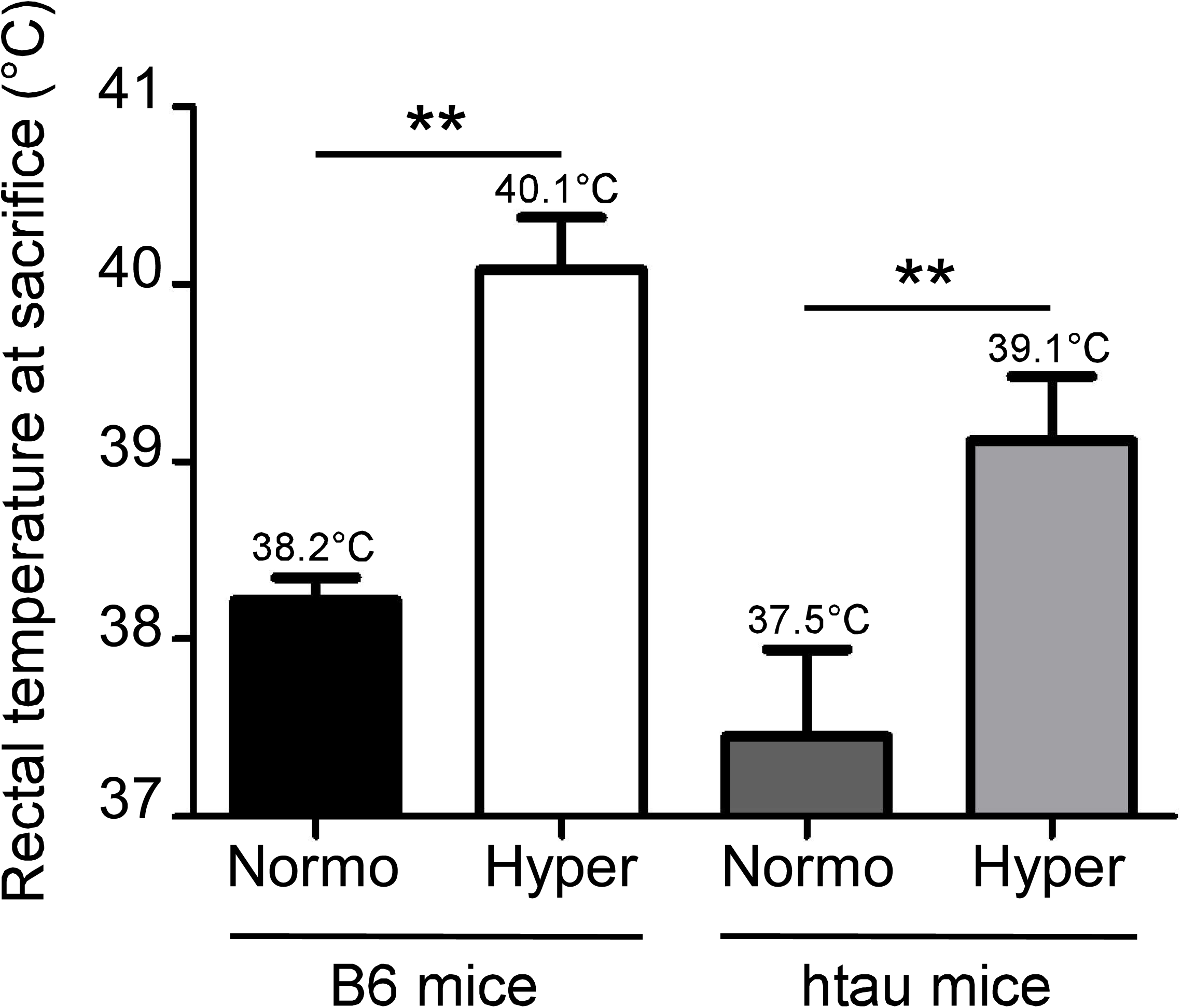
Mild hyperthermia increases the rectal temperature of B6 (n=12) and hTau mice (n=11). Temperatures of the mice kept at room temperature (Normo) or exposed at 42°C for 45min (Hyper). For both genotypes the mean rectal temperature of the mice housed at room temperature was significantly different from that of the mice housed at 42°C. The numbers written above each bar indicate the average rectal temperature measured. Bilateral T-test, p <0.0002 (***).

**Figure 2:**
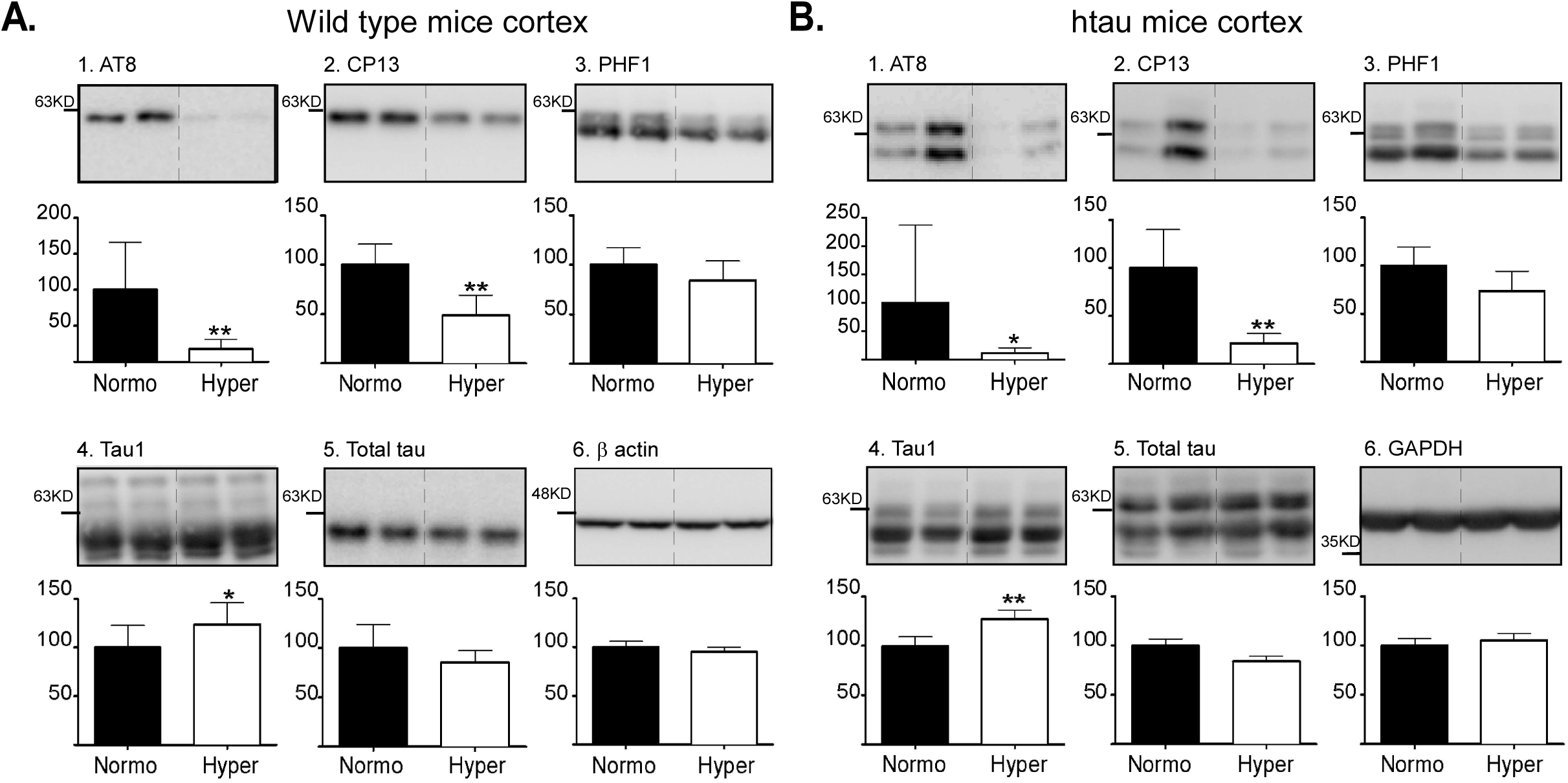
Tau phosphorylation in the cortex of B6 mouse (A; n=12) and hTau mouse (B; n=11) in condition of normothermia (Normo) and of mild hyperthermia (Hyper). The following antibodies were probed: (1) AT8, (2) CP13, (3) PHF1, (4) Tau1, (5) total tau and (6) b actin. Data are expressed as the percentage of signal obtained at normothermia (y-axis). In B6 mice, Tau was significantly dephosphorylated at all epitopes tested. In hTau mice, Tau was significantly dephosphorylated at all epitopes tested except PHF1. Two tailed Mann-Whitney Testing, results are given for p<0.009, a < 0,01 (**) and p <0.033, a < 0,05 (*).

**Figure 3:**
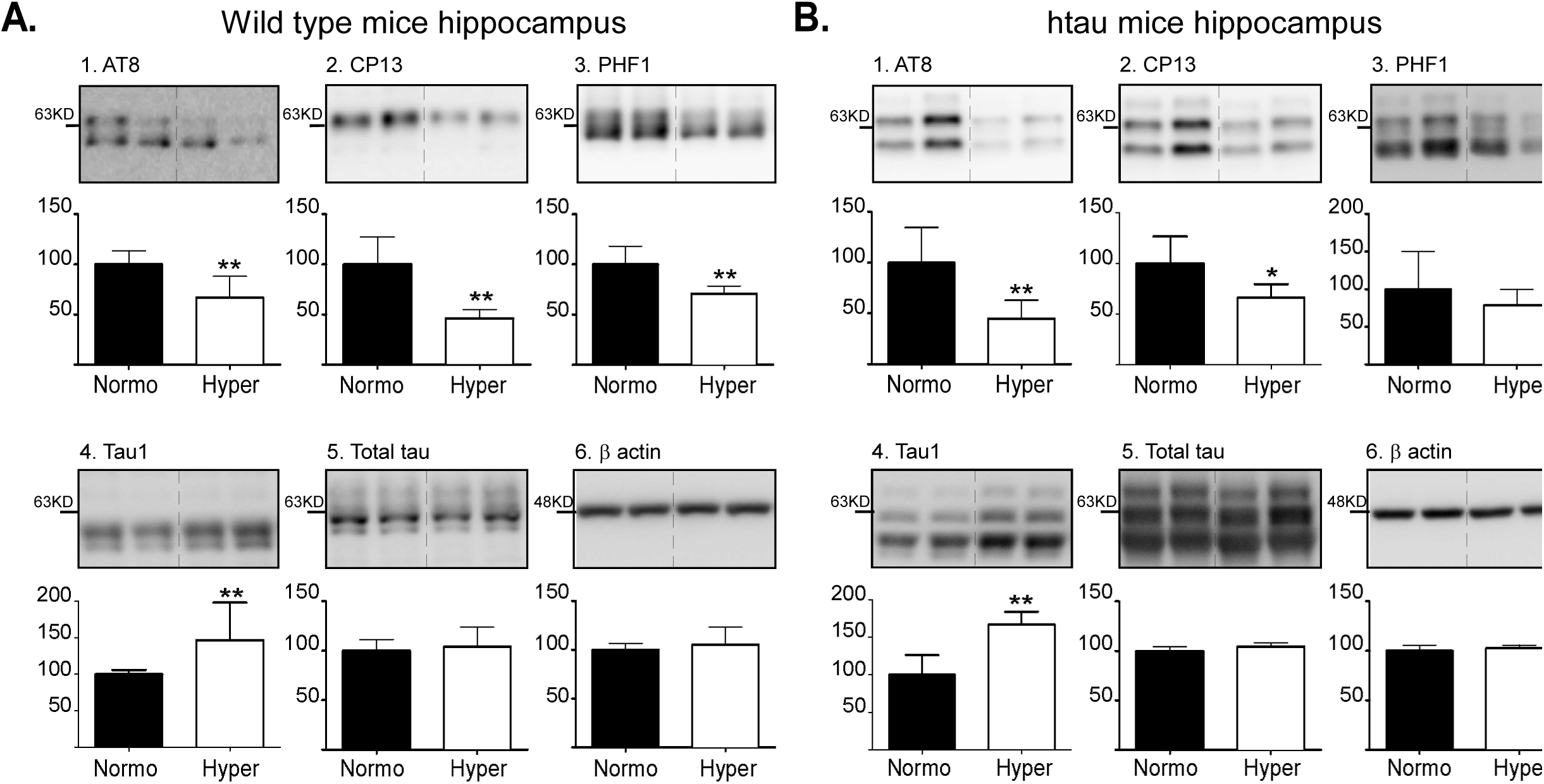
Tau phosphorylation in the hippocampus of B6 mouse (A; n=12) and hTau mouse (B; n=11) in condition of normothermia (Normo) and of mild hyperthermia (Hyper). The following antibodies were probed: (1) AT8, (2) CP13, (3) PHF1, (4) Tau1, (5) total tau and (6) b actin. Data are expressed as the percentage of signal obtained at normothermia (y-axis). In B6 mouse, Tau was dephophorylated at all epitopes tested. In hTau mouse, Tau was dephosphorylated at AT8 and CP13 but not at AT180 and PHF1. Two tailed Mann-Whitney Testing, results are given for p<0.009, a < 0,01 (**) and p<0.031, a < 0,05 (*).

**Figure 4:**
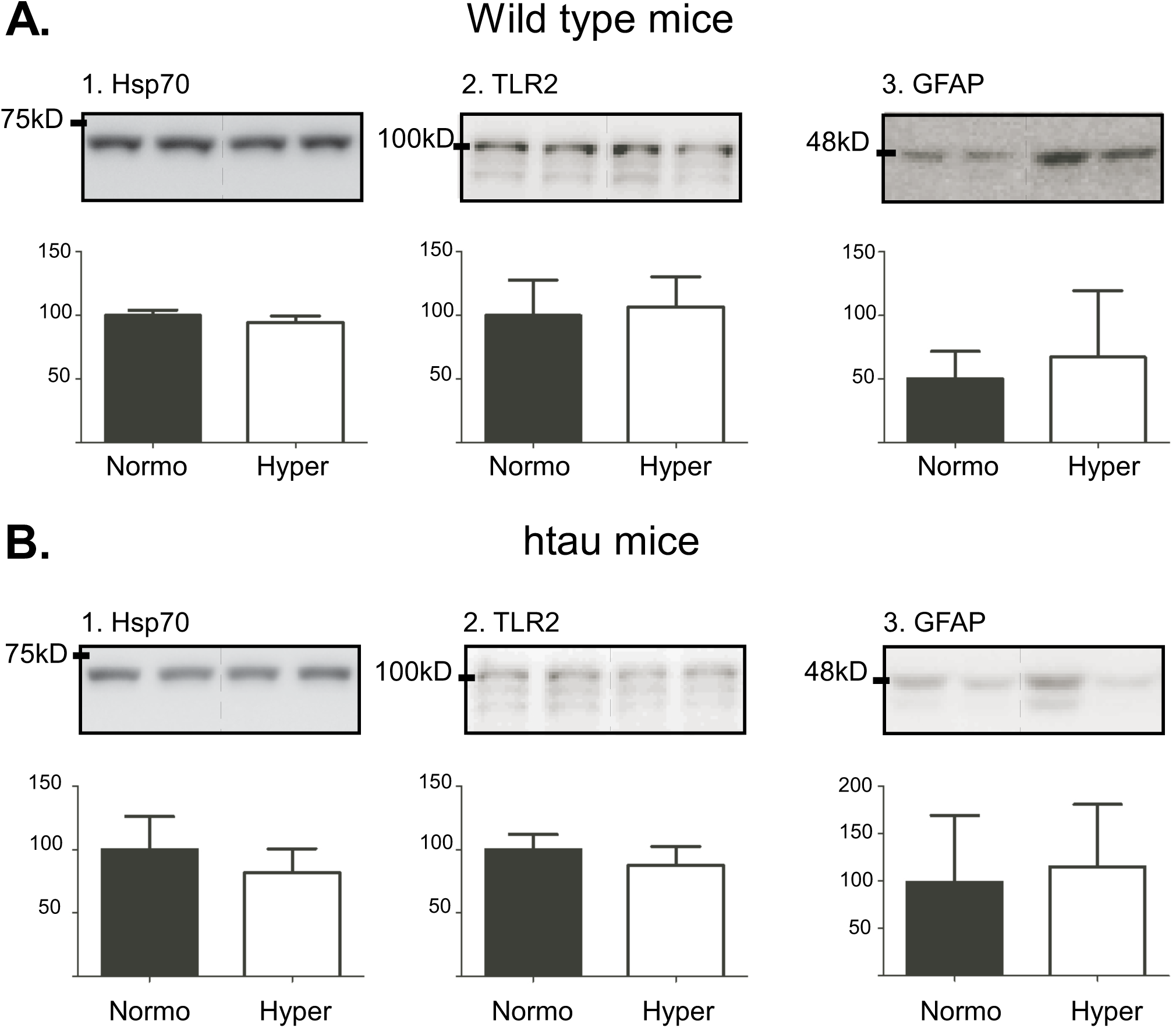
Effect of mild hyperthermia on inflammation on the cortex of B6 mice (A) and hTau mice (B) (n=6 in each temperature except for GFAP: n=12). The following antibodies were probed: (1) HSP70, (2) TLR2 and (3) GFAP. Data are expressed as the percentage of signal obtained at normothermia (y-axis). Mild hyperthermia didn’t modify expression of markers of inflammation. (Two tailed Mann-Witney Testing).

**Figure 5:**
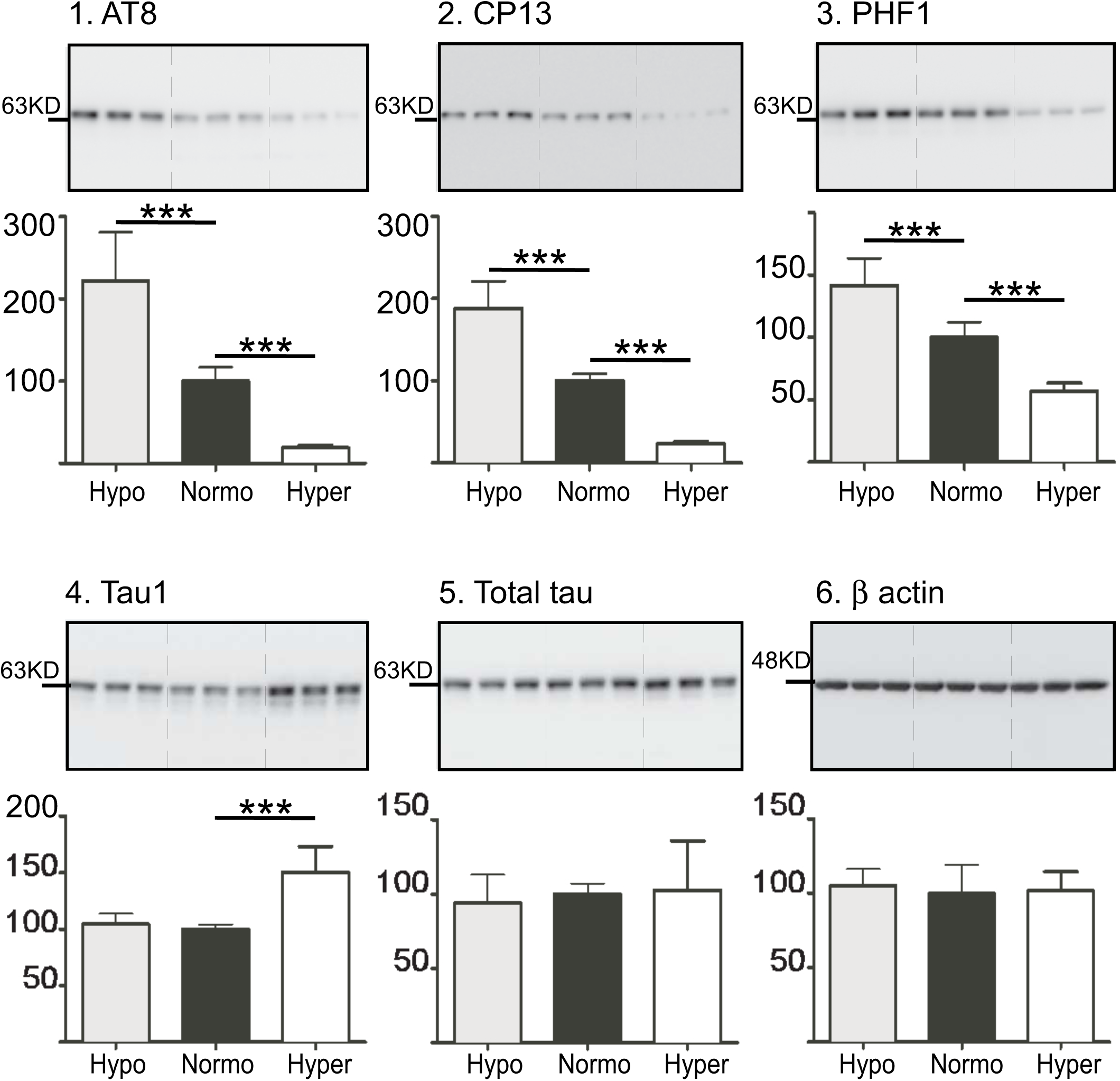
Tau phosphorylation in SHSY5Y-1N3R (n=9/group) in condition of hypothermia (34°C), normothermia (Normo, 37°C) and of mild hyperthermia (40°C). The following antibodies were probed: (1) AT8, (2) CP13, (3) PHF1, (4) Tau1, (5) total tau and (6) b actin. Data are expressed as the percentage of signal obtained at normothermia (y-axis). De-phosphorylation of Tau was observed at all epitopes upon hyperthermia. Two tailed Mann-Whitney Testing, results are given for p<0.0005 and a<0.01 (***).

## Results

### Mild hyperthermia lowers tau phosphorylation in wild type and hTau mice

We first wanted to assess whether mild and transient hyperthermia could affect tau phosphorylation *in vivo* in mice. We thus explored the effects of one hour of hyperthermia on tau phosphorylation in C57Bl6 mice and in a mouse model of human tauopathy, hTau mice. Both B6 (Ctrl: 38.22 ± 0.13°C; Hyper: 40.08 ± 0.29°C), and hTau (Ctrl: 37.45 ± 0.49°C; Hyper: 39.12 ± 0.36°C) mice exposed to heat displayed mild hyperthermia at sacrifice compared to the control mice housed at room temperature **(Fig.1)**. Tau was dephosphorylated at the AT8, CP13, and Tau1 epitopes in the cortex of C57B6 and hTau mice **(Fig.2)**. We next wanted to investigate whether the dephosphorylation of Tau was the same in the hippocampus, one of the first regions to display tau pathology in AD (Braak et al. 2006). Again, we observed significant hippocampal dephosphorylation of tau protein at epitopes AT8, CP13, PHF1 and Tau1 in C57Bl6 mice **(Fig3.A)** and at the AT8, CP13 and Tau1 epitopes of hTau mice **(Fig3.B)**. However, the treatment did not impact tau aggregation in hTau mice as evaluated by sarkosyl extraction (data not shown). Overall our data demonstrate that mild hyperthermia can lead to extensive tau dephosphorylation at multiple epitopes.

### Mild hyperthermia does not trigger heat-shock response or subsequent inflammatory

We next wanted to assess whether mild hyperthermia had the potency to trigger a heat-shock response and subsequent inflammation. Indeed, heat-shock can induce markers of heat-shock and inflammation such as HSP70 (Heat-Shock Protein 70), TLR2 (Toll Like Receptor 2) and GFAP (Glial Fibrillary Acidic Protein) (Brown 1983, Miller et al. 1987, Schiaffonati et al. 2001, Belay and Brown 2003, Hayward and Lee 2014, Lee et al. 2015), that can modulate tau pathology (Bolos et al. 2017). Our results show that none of the three markers were increased in C57Bl6 or hTau mice (**Fig.4**), suggesting that at 39-40°C hyperthermia was not enough to induce heat shock or an inflammatory responses in our experimental conditions. In turn, these results suggest that neither heat-shock nor inflammatory responses are responsible for tau dephosphorylation in our mice.

### Mild hyperthermia lowers tau phosphorylation in neuroblastoma cells

To isolate temperature from confounding factors such as blood pressure, we explored the direct effects of mild hyperthermia (37°C vs 40°C) on the phosphorylation of human tau protein in SH-SY5Y 3R-tau cells. Strikingly, we observed that tau protein was strongly dephosphorylated at the same epitopes as seen in mice (AT8, CP13, PHF1, and Tau1 (tau1 recognizes dephosphorylated epitopes) **(Fig.5)**. We also used a positive control for tau hyperphosphorylation by exposing cells to hypothermia. As previously described (Bretteville, Marcouiller et al. 2012), tau protein was hyper phosphorylated at 34°C at all epitopes tested in comparison to 37°C. These results suggest that a mild increase in temperature alone is sufficient to induce neuronal tau dephosphorylation.

### Mild hyperthermia inhibits JNK activation

We next aimed to better understand the underlying molecular mechanisms of tau dephosphorylation. Since tau phosphorylation is a balance between the activation state of tau kinases and phosphatases (Wang et al. 2007, Martin et al. 2013, Iqbal et al. 2016), we speculated that hyperthermia could induce a imbalance of the system towards kinases inhibition and/or phosphatases activation. We thus analyzed the activation state (phosphorylation) of some of the major kinases involved in the regulation of tau phosphorylation: JNK (c-Jun N-terminal Kinase), GSK-3b (Glycogen Synthase Kinase 3b), AMPK (Adenosine Mono Phosphate activated protein Kinase), PKA (Protein Kinase A), ERK (Extra-cellular Responsive Kinase), CAMKII (Calcium-Calmodulin dependent protein Kinase II), and AKT (Protein Kinase B) (Wang, Grundke-Iqbal et al. 2007, Cai et al. 2012, Martin, Latypova et al. 2013, Iqbal, Liu et al. 2016). We also analyzed the activation state of PP2A (methylated PP2A), a phosphatase involved in AD (Wang, Grundke-Iqbal et al. 2007, Martin, Latypova et al. 2013, Iqbal, Liu et al. 2016) and responsible of 70% of tau phosphatase activity in the human brain (Liu et al. 2005). We observed that JNK was inactivated in cortex of C57Bl6 and hTau mice by heat exposure **(Fig.6)**, while there was no significant changes in the activation state of all the other kinases or PP2A. Importantly, the antibodies directed against the total forms of these enzymes did not show any change (data not shown). Our results show that JNK is partially inactivated during hyperthermia.

**Figure 6:**
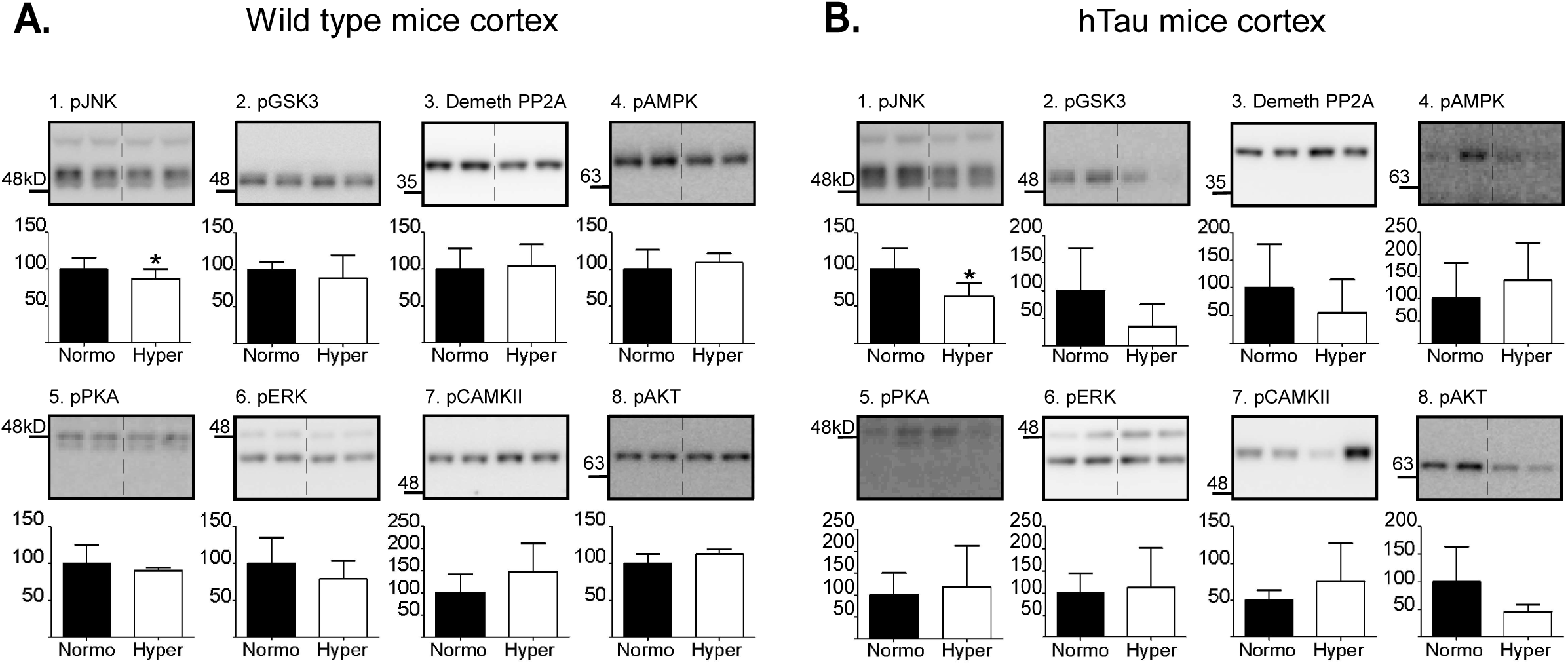
Activation of kinases and phosphatases of tau in the cortex of B6 mice (n=12) (A) and hTau mice (n=11) (B) in condition of normothermia (Normo) and of mild hyperthermia (Hyper). The following antibodies were probed: (1) pJNK, (2) pGSK3b, (3) demethylated PP2A, (4) pAMPK, (5) pPKA, (6) pERK, (7) pCAMKII, (8) pAKT. Data are expressed as the percentage of signal obtained at normothermia (y-axis). c-Jun N terminal Kinase (JNK) was significantly inactivated with hyperthermia (in both B6 and hTau mice) while other kinases and phosphatases were not. Two tailed Mann-Whitney Testing, results are given for p<0.047, a < 0,05 (*).

### Hyperthermia results in an imbalance between tau phosphatases and kinases activities

While, we show that JNK is partially inactivated, it is not sufficient to explain the extent of tau dephosphorylation observed. From our previous work, we know that lower temperatures can directly and differentially affect the activities of phosphatases and kinases, resulting in tau hyper-phosphorylation (Planel, Miyasaka et al. 2004), we thus explored whether higher temperature could also modulate kinases and phosphatases activities. We assessed PP2A activity (a major phosphatase of tau representing 70% of tau phosphatases activity (Liu, Grundke-Iqbal et al. 2005)) and total tau kinases activity in brain lysates, when exposed to high temperatures (40°C and 42°C) in comparison to controls at 37°C. Interestingly, we observed an increase in PP2A activity at 40°C and 42°C **(Fig.7.A)** while kinases activity at the same temperatures was decreased **(Fig.7.B)**. Thus increasing body temperature can promote an imbalance between kinases and phosphatases favoring tau dephosphorylation.

**Figure 7:**
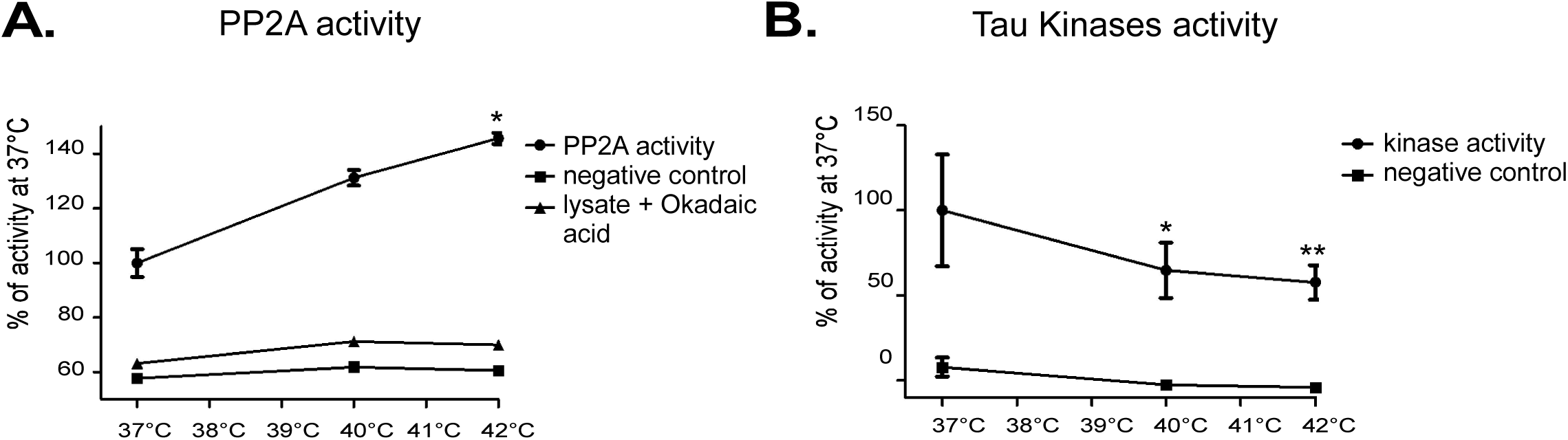
Effect of temperature on PP2A activity (A) and on Kinases activity (B). **(A)**. Phosphatase activity observed by green malachite colorimetry in total protein extracted from cortex of a B6 mouse at different temperatures (n=3 in each temperature). The activity is expressed as a percentage of activity at 37°C. Phosphatase activity increased significantly with temperature. Kruskall Wallis test followed by Dunn’s post hoc test for pair-wise sample comparison. Significant results are given for p<0.05 (*). **(B)**. Effect of temperatures on the activity of kinases *in vitro* (n=6 in each group). The activity is expressed as a percentage of activity at 37°C. Kinase activity decreased significantly with temperature. Signal is normalized to total tau. One Way ANOVA with Holm Sidak’s multiple comparison test, p< 0,05 (*), p<0.01 (**).

### Topical application of menthol increases body temperature phosphorylation and lowers tau

As sauna bathing is not available to most people, we thought of an alternative manner to increase body temperature. Menthol is able to increase body temperature by acting on TRPM8 channels (Tajino, Matsumura et al. 2007). We found that topical application of menthol increased body temperature by 1.3°C 2 hours after application **(Fig.8.A**). This resulted in a decrease of tau phosphorylation in the cortex **(Fig.8.B)**:. AT8 (−70%), PHF1 (−25%) and Tau1 (+25% as tau 1 recognizes dephosphorylated tau). As menthol targets TRPM8 and as mice lacking TRPM8 have an enhanced clearance of insulin, a parameter known to potentially modulate tau phosphorylation (Lee et al. 2013, Lauretti et al. 2017) we evaluated whether glycaemia was affected by menthol exposition. Our result show that glycaemia was unaffected by menthol topic application **(Fig.8.A**). Altogether, these results demonstrate that increasing body temperature pharmacologically by menthol treatment lowers tau phosphorylation in the brain.

**Figure 8:**
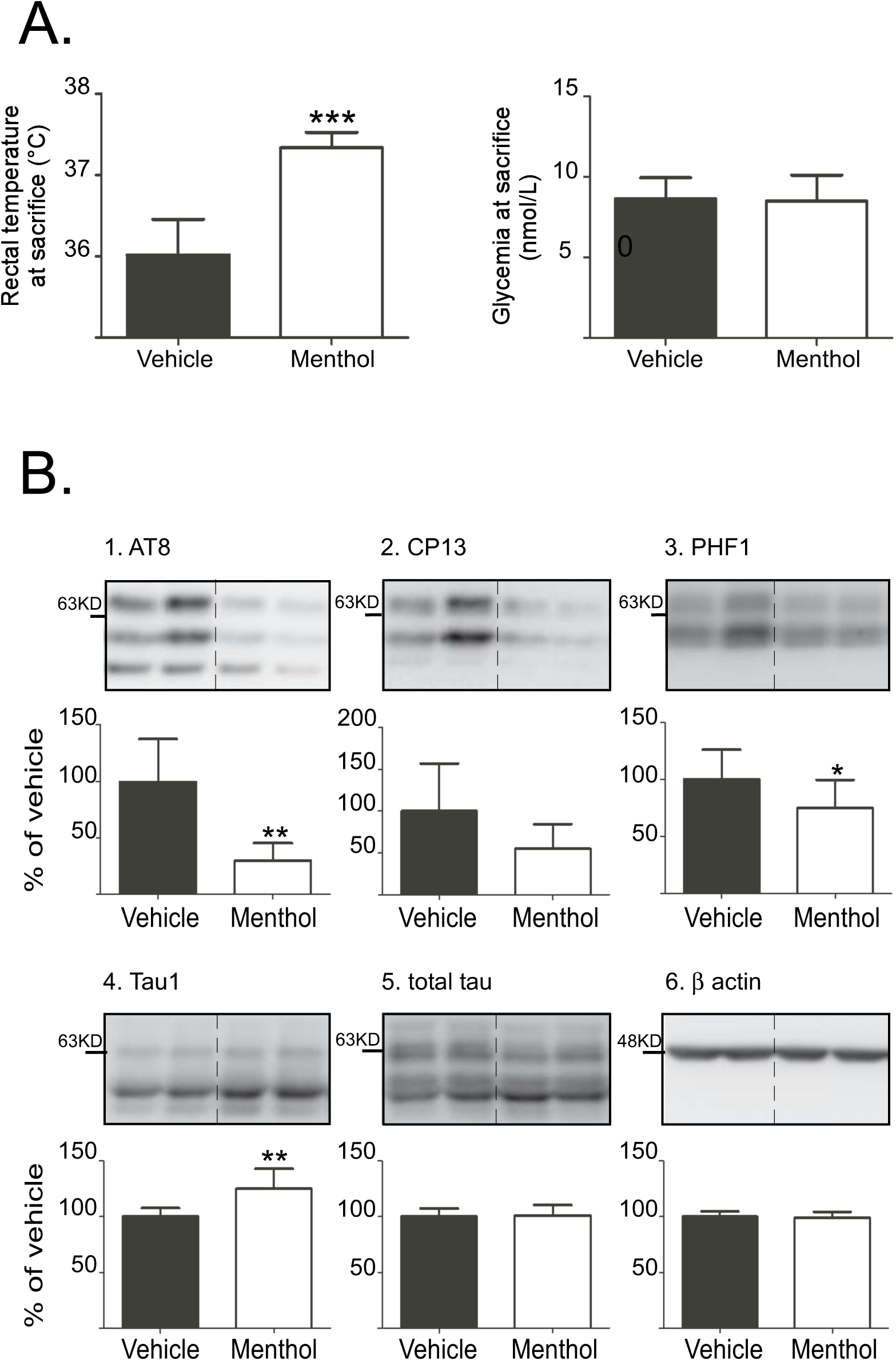
Topical application of menthol is able to increase rectal temperature (A) and to decrease phosphorylation of tau in the cortex of hTau mice. **(A)**. Two hours after ectopic exposition to menthol, rectal temperature and glycaemia were assessed (vehicle: n=6; menthol: n=9). Rectal temperature was significantly increased under ectopic application of menthol although glycaemia was unaffected. Unpaired t-test with welch correction, p=0.0005, (***) **(B)**. Phosphorylation of tau in the cortex of hTau mice 2 hours after one ectopic application of menthol. Tau is significantly dephosphorylated at AT8, PHF1, and Tau1. Unpaired t-test with welch correction, p<0.01 (**), and p<0.05 (*).

### Heat exposure or menthol treatment increase body temperature without subsequent hypothermia

To explore the impact of heat or menthol treatment on core body temperature, we used telemetry in mice exposed to 42°C for 45min **(Fig.9.A)** and in 10% menthol treated mice **(Fig.9.B)** compared to their respective controls. Increased body temperature peaked 60 min after the beginning of heat exposure (+3.3°C, ±0.27 p<0.0001) and was significant up to 76 min after (+1.3°C, ±0.17, p=0.0004) in heat treated mice. The temperature of menthol treated mice was highest (+2°C, ±0.1 p<0.0001) 2 h 40 min after exposure to menthol and the last significant difference (+0.8°C, ±0.21, p<0.01) was detected after 8 h 30 min. Heat-shock in rodents can be followed by hypothermia and subsequent tau hyperphosphorylation (Papasozomenos 1996, Leon et al. 2005). Here, mild hyperthermia induced by either heat- or menthol-treated mice did not lead to hypothermia.

**Figure 9:**
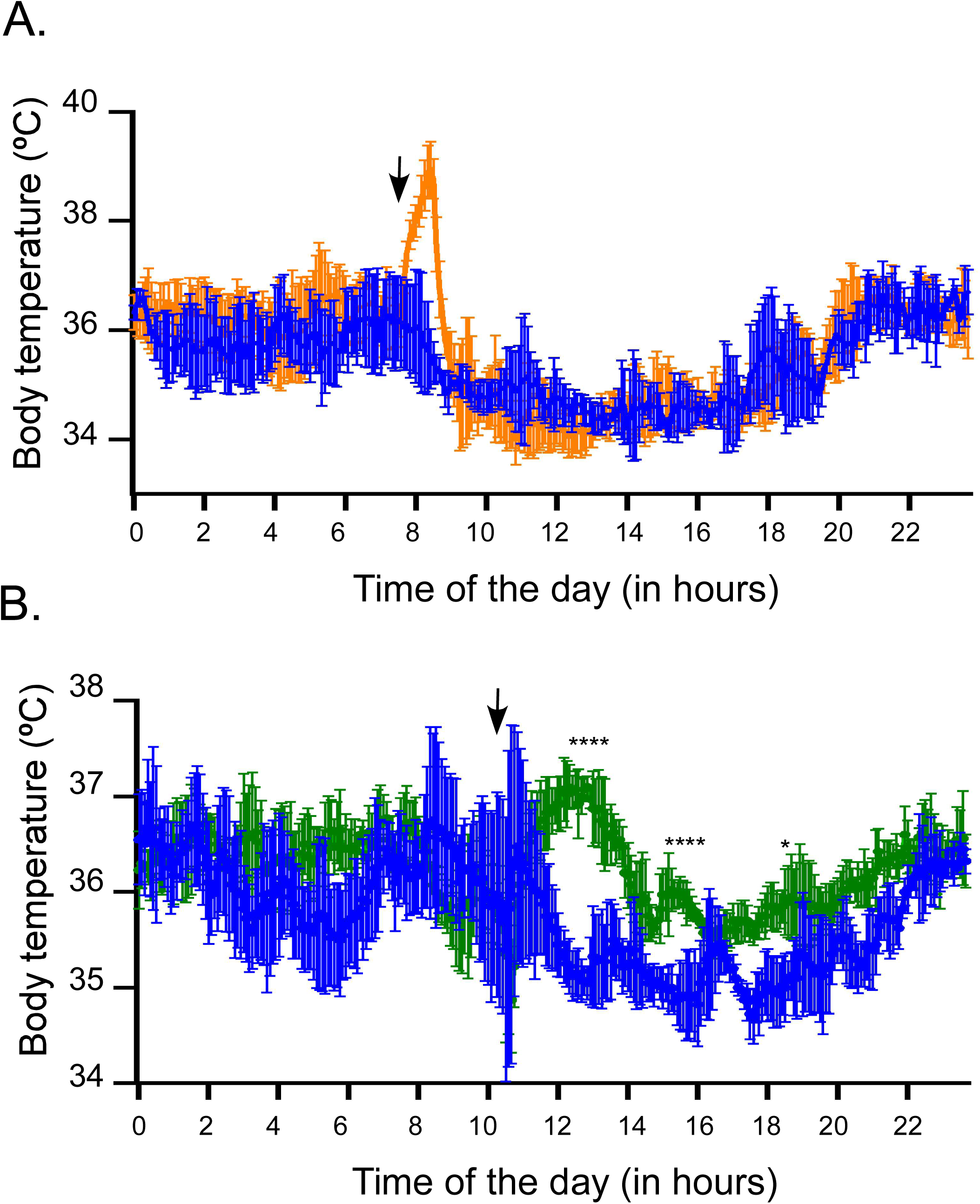
Heat treatment (A) and menthol treatment (B) increase significantly body temperature but do not induce hypothermia after treatment. Core body temperature was assessed in B6 mice with probes surgically implanted in the abdomen of the animals **(A)**. The orange line represents the mean body temperature of the animals exposed to heat (42°C, 45min, n=4). Blue line gives the mean of body temperature of animals housed at room temperature (23°C, n=4) during the experiment. Time is given as time of the day. During the heat exposure, the body temperature of heat exposed animals was significantly higher than body temperature of animals housed at room temperature at 8h08min (+2.5°C, p=0.0147), at 8h27min (3.4°C, p<0.0001) and at 8h46min (+1.3°C, p=0.0004). Unpaired t-test with welch correction. **(B)** The green line represents the mean body temperature of the animals exposed to 10% menthol (Topic exposition, n=4). Blue line gives the mean of body temperature of animals exposed to vehicle (Topic exposition, n=4). Time is given as time of the day. During the menthol exposure, the body temperature of mint exposed animals was significantly higher than body temperature of animals exposed to vehicle at 12h40min (+2°C, p<0.0001, ****), 15h14min (+1.25°C, p<0.0001, ****) and 18h28min (0.8°C, p=0.014, *). Unpaired t-test with welch correction.

## Discussion

In this study, we investigated for the first time the effects of mild hyperthermia on tau phosphorylation *in vivo* and *in vitro*. We report that mild hyperthermia lowers tau phosphorylation in both non-transgenic and hTau mice, and in neuron-like cells expressing human tau, We also observed that these changes were probably the consequence of an imbalance between phosphatase and kinase activities favoring tau dephosphorylation.

Tau hyperphosphorylation and aggregation are associated with the progression of AD (Simic et al. 2016). Here, we showed that mild hyperthermia led to dephosphorylation of tau at epitopes that are known to be hyper-phosphorylated in AD: AT8, CP13, Tau1 and PHF1 (Simic, Babic Leko et al. 2016, Tremblay et al. 2017). The AT8 antibody is used for AD brain staging in a more sensitive manner than silver staining (Braak, Alafuzoff et al. 2006). Moreover, phosphorylation at both AT8 (Ser 202 and Thr205) and CP13 (Ser202) epitopes can enhance tau aggregation (Rankin et al. 2005), and AT8 phosphorylation can enhance the phosphorylation of other epitopes (Bertrand et al. 2010). Importantly, Ser202 is an early marker of AD (Su et al. 1994) and is suggested as a major contributor of neurofibrillary tangles formation (NFT) (Han et al. 2009). Hyperphosphorylation at PHF1 epitope is one of the earliest event that occurs in AD brain (Mondragon-Rodriguez et al. 2014). Tau hyperphosphorylation can lead to its aggregation (Alonso, Grundke-Iqbal et al. 1996), so we initially thought that hyperthermia would decrease tau aggregates. However, acute mild hyperthermia did not reduce tau aggregation in hTau mice. As the sauna bathing study showed that high frequency sauna users were at lower risk to develop AD (Laukkanen T.S.K 2016) our acute exposure study might not have been sufficient to reduce tau aggregation. It would thus be interesting in the future to test whether chronic or repeated exposition to mild hyperthermia decreases tau aggregation. Altogether our results demonstrate that hyperthermia reduces tau phosphorylation at key pathological epitopes and suggest that it could be beneficial by reducing tau pathology.

While the effects of heat shock on tau phosphorylation has been reported, the response to mild physiological hyperthermia has not been investigated. For instance, Papasozomenos *et al*. demonstrated that upon 45°C heat-shock in rat fetal cerebral explants, tau protein is rapidly dephosphorylated (Papasozomenos and Su 1995). Sultan *et al*. observed also that upon heat shock in mouse primary embryonic culture (44°C), tau is dephosphorylated (Sultan et al. 2011). Heat-shock induces HSPs expression and inflammatory responses (Leon 2007, Laurent et al. 2018), which can both modulate tau phosphorylation (Kirby et al. 1994). Here, we observed that inflammatory and heat-shock processes were not induced in our animals during mild hyperthermia, in contrast to previous published work on heat-shock (Papasozomenos and Su 1995, Papasozomenos 1996, Sultan, Nesslany et al. 2011). Importantly, heat-shock in rodents (42-43°C body temperature) is followed by several hours of hypothermia (Leon, DuBose et al. 2005), and can lead to tau hyperphosphorylation (Papasozomenos 1996). As we and others have shown that hypothermia can increase tau phosphorylation and aggregation (Planel, Miyasaka et al. 2004, Planel, Richter et al. 2007, Maurin et al. 2014, Carrettiero et al. 2015, de Paula et al. 2016, Vandal, White et al. 2016, Almeida and Carrettiero 2018), it was important to ascertain that our experimental paradigm did not induce subsequent hypothermia. Our telemetry data demonstrate that it was not the case. Thus, our results suggest that mild hyperthermia *per se* is capable of lowering tau phosphorylation without inducing inflammatory, heat-shock responses, or hypothermia.

We next explored the molecular mechanisms underlying tau dephosphorylation and found that the effect of mild hyperthermia on tau phosphorylation was probably mediated through a direct effect of temperature on kinases and phosphatases activities resulting in tau de-phosphorylation. More specifically, total tau kinase activity was reduced during heating, a result in line with those of Shavanas and Papasozomenos, who observed a suppression of nearly all kinases activities during heat shock (Shanavas and Papasozomenos 2000). We also report that mild hyperthermia lowered JNK activation. This observation is concordant with two previous studies. One showing that in adult rat cerebellum, JNK1 and JNK3 are deactivated by heat-shock (Schiaffonati, Maroni et al. 2001) and the other showed that global JNK activity decreases at high temperatures (Shanavas and Papasozomenos 2000). We also observed an increase in PP2A activity, which is consistent with the prevention of tau dephosphorylation by PP2A inhibitors during heat-shock (Papasozomenos and Su 1995). PP2A activity is down regulated in AD concomitant with tau hyper-phosphorylation (Sontag and Sontag 2014). Furthermore, the activation of PP2A has already been shown to reverse tau hyperphosphorylation (Chohan et al. 2006, van Eersel et al. 2010, Luo et al. 2013, Song et al. 2014, Yang et al. 2016), and to have beneficial effects on memory, motor performance and neurodegeneration in animal models of the disease (van Eersel, Ke et al. 2010). Since our results show that mild hyperthemia can increase PP2A activity and reduces abnormal phosphorylation of tau, our results suggest that heat exposure could be beneficial for targeting tau pathology through activation of PP2A.

Sauna bathing decreases blood pressure in hypertensive patients (Winterfeld et al. 1993, Hannuksela and Ellahham 2001) and hypertension in mid-life is associated with a higher risk of developing AD (Glodzik et al. 2014, Edwards Iii et al. 2019). Interestingly, inducing hypertension in mice models of AD, is able to increase tau phosphorylation in the brain (Shih et al. 2018). Sauna bathing could therefore decrease phosphorylation of tau through modulations in blood pressure. However, we observed that tau was dephosphorylated in neuronal cells exposed to mild hyperthermia, suggesting that mild hyperthermia can lead to neuronal tau dephosphorylation, independently of potential effects of blood pressure.

As sauna bathing is not readily available to most people, we thought of a therapeutic way to increase body temperature using topical menthol treatment, which results in TRPM8 activation and promotes thermogenesis and temperature increase in mice and humans (Tajino, Matsumura et al. 2007, Ma et al. 2012, Gillis, Weston et al. 2015). Our telemetry results show that menthol induced a transient increase in body temperature and consequent tau dephosphorylation. Menthol is an organic compound made synthetically or obtained from the oils of mint plants such as peppermint (*Mentha* × *piperita)*. Peppermint oil has been used for centuries as a treatment for gastrointestinal ailments. Its effects include smooth muscle relaxation, visceral sensitivity modulation; anti-microbial effects, anti-inflammatory activity and it may play a role in modulation of psychosocial distress (Chumpitazi et al. 2018). Menthol has also been shown to protect from to β-amyloid peptide induced cognitive deficits in mice (Bhadania et al. 2012), to protect from glutamatergic excitotoxicity (Pezzoli et al. 2014), and to be a non-competitive inhibitor of serotonergic receptors (Ashoor et al. 2013). Moreover, menthol derivatives were shown to inhibit acetylcholinesterase and butyrylcholinesterase, making them drug candidates for Alzheimer’s disease (Daryadel et al. 2018), especially as menthol is known to cross the blood brain barrier (Pan et al. 2012). However, the effects of menthol on tau pathogenesis had not been tested until now. Together, previous publications and our present work suggest that menthol has pleiotropic effects on many features of AD and could be a promising therapy for this disease.

TRPM8 is expressed in somatosensory neurons that innervate peripheral tissues and mediates thermoregulatory responses upon activation (Bhadania, Joshi et al. 2012, Ashoor, Nordman et al. 2013, Pezzoli, Elhamdani et al. 2014, Daryadel, Atmaca et al. 2018). Aside from its effects in the nervous system, TRPM8 is also expressed in white adipose tissue and its specific activation by menthol mediates acquisition of “brown like phenotype” (“browning”) (Rossato et al. 2014). Indeed, in primary culture of white adipocytes (Rossato, Granzotto et al. 2014) and *in vivo* (Ma, Yu et al. 2012), menthol specific stimulation of TRPM8 increases expression of UCP1 (uncoupling protein 1) a protein responsible for transferring back proton from the inner membrane space to the mitochondrial matrix and resulting in heat production. Consistent with these data, menthol ingestion has also been shown to promote higher temperatures in humans, although to a lesser extent than topical application, probably because of high metabolism (Valente et al. 2015). Thus, *per se* delivery of menthol or derivatives tailored for slower degradation could stimulate thermogenesis *via* white adipose browning and be a simpler treatment for AD.

Old age is the most important risk factor for AD (Harman 2002, Ritchie and Lovestone 2002), and older adults have a tendency to display cold intolerance, like sweaters in summertime, or hothouse conditions in winter (Holtzman and Simon 2000). Indeed, elderly humans regulate core temperature less efficiently than younger adults in hot or cold environments (Horwitz 2001), and many of them have lower body temperatures (Fox et al. 1973, Keilson 1985, Collins 1992). They are prone to hypothermia for a variety of reasons including decreased basal metabolism, decreased thermogenesis, less intense shivering, decreased muscle mass that generate fewer calories, loss of body fat, decreased sensitivity to cold and changes in temperatures, and abnormal vasoconstrictor responses to cold (Brody 1994). We were the first to observe that reduced body temperature leads to tau hyperphosphorylation both *in vivo* and *in vitro* (Planel, Richter et al. 2007), and can promote long-term increase in tau pathology in mice models of AD (Planel et al. 2009). These pre-clinical data, combined with the above-mentioned clinical evidence, suggest that an increased incidence of hypothermia during aging may contribute to tau pathogenesis (Whittington, Papon et al. 2010, Almeida and Carrettiero 2018). Interestingly, restoring thermoneutrality or promoting thermogenesis in animal models of AD decreased amyloid deposits or prevented tau hyperphosphorylation after a cold challenge (Vandal, White et al. 2016, Tournissac et al. 2019). Here, we show that promoting higher body temperature by higher ambient temperature or pharmacologically promoting thermogenesis can decrease tau phosphorylation. Our findings might represent a way to counteract development of tau hyperphosphorylation in elderly patients with thermoregulatory deficits.

In conclusion, we show for the first time that mild acute hyperthermia is able to decrease tau phosphorylation in mice by activating PP2A and inhibiting specific kinases. Our discovery highlights the fact that controlling body temperature might represent a potential therapeutic tool in regard of AD and is sustained by the fact that sauna bathing studies in humans have important benefit effects in AD (Laukkanen T.S.K 2016). This potential therapy has the potential advantage to reverse tau phosphorylation in a physiological manner that might be easily applicable in humans. Further evaluation of chronic or repeated exposure to mild hyperthermia, or longer menthol treatment in mouse models of AD should be done to assess its benefit effect on the progression of the disease.

